# Disconnect between *in vitro* and *in vivo* efficacy of the MPS1 inhibitor NTRC 0066-0 against glioblastoma

**DOI:** 10.1101/2024.11.04.621827

**Authors:** Mark C. de Gooijer, Ping Zhang, Levi C.M. Buil, Ceren H. Çitirikkaya, Hilal Çolakoğlu, Ana Rita R. Maia, Irena Bočkaj, Margarita Espitia-Ballestas, Laura E. Kuil, Jos H. Beijnen, Olaf van Tellingen

**Author notes:** **To whom correspondence should be addressed**: O. van Tellingen; Division of Pharmacology, room H3.014; Plesmanlaan 121, 1066CX Amsterdam, The Netherlands. **Statements and declarations**. **Competing Interests:** The authors declare no competing interests. **Funding:** This work was funded by a grant from Stichting STOPHersentumoren.nl to OvT. **Data availability statement:** all data will be uploaded to a repository prior to final publication of the manuscript. **Author contributions:** JHB, OvT, and MCdG conceived the study. MCdG, PZ, LCMB, CHÇ, HÇ, ARRM, IB, MEB and LEK collected and analyzed data. JHB and OvT, supervised the study. OvT acquired funding for the study. MCdG and OvT wrote the manuscript with input from all authors.

## Abstract

**Purpose:** Glioblastoma (GBM) is the most common adult primary brain tumor for which new therapeutic strategies are desperately needed. Monopolar spindle 1 (MPS1) is a mitotic kinase that plays a pivotal role in the spindle assembly checkpoint (SAC). GBM appears to be dependent on SAC fidelity, as MPS1 is overexpressed in many GBM patients. Thus, inhibiting MPS1 seems a viable therapeutic strategy to enhance mitotic cell death by attenuating SAC fidelity. NTRC 0066-0 is an MPS1 inhibitor that combines low nanomolar potency with a relatively long on-target residence time.

**Methods:** We here investigate the potential of NTRC 0066-0 as monotherapy and in combination with chemo-radiation for treatment of GBM using various *in vitro* and orthotopic *in vivo* models.

**Results:** We show that NTRC 0066-0 efficiently induces GBM cell death *in vitro*, following continuous exposure with IC_50_s in the low nanomolar range. In contrast to previous reports of studies with other MPS1 inhibitors, we did not observe synergy *in vitro* with anti-microtubule drugs, such as docetaxel and vincristine. We demonstrate that NTRC 0066-0 has a high brain penetration, despite being a substrate of the efflux transporter P-glycoprotein. However, even when using recipient *Abcb1a/b;Abcg2^-/-^* mice with superior brain penetration and administering NTRC 0066-0 using a dose-dense regimen, we did not observe antitumor efficacy against an orthotopic GBM mouse model, neither as monotherapy nor in combination with standard-of-care temozolomide chemotherapy and radiotherapy.

**Conclusion:** These data indicate that GBM is probably not a suitable indication for developing MPS1 inhibitors.

## 1. INTRODUCTION

Glioblastoma (GBM) is the most common adult primary brain tumor and has a dismal median survival of only 15 months [1]. Since the addition of the alkylating agent temozolomide to the standard of care in 2005, no other drugs have significantly improved the survival of glioblastoma patients [1]. New therapeutic options are thus needed, and several avenues are currently being explored. One of these is targeting monopolar spindle 1 (MPS1; HUGO name, TTK), a serine/threonine kinase that is an integral part of the spindle assembly checkpoint (SAC) [2]. MPS1 safeguards genomic integrity throughout mitosis by inducing and maintaining a mitotic arrest when chromosomes are improperly connected to the mitotic spindle, thereby preventing missegregation events that may result in chromosomal instability (CIN) [3]. GBMs seem to depend more heavily on the SAC, as several SAC components are overexpressed in GBM compared to low-grade gliomas and healthy brain tissue, including budding uninhibited by benomyl 1 (BUB1) and MPS1 [4, 5]. As replication stress increases the chance of missegregations [6], the upregulation of critical SAC components may reflect a strategy of GBMs to cope with increased intrinsic levels of DNA damage. Attenuating SAC signaling by inhibiting MPS1 might therefore force GBM cells to continue division without proper chromosome arrangement, resulting in CIN and ultimately cell death. Importantly, GBM cells might even depend more heavily on the SAC following exposure to the standard of care chemo-radiotherapy, as this induces genome-wide DNA damage.

Several MPS1 inhibitors have been developed to date [7–10], and one compound, MPS1-IN-3, has even demonstrated modest efficacy in a preclinical model of GBM when combined with the microtubule poison vincristine [5]. While these early results with MPS1-IN-3 were encouraging, it has not been taken into further clinical development. Notably, the intrinsic potency of this MPS1 inhibitor is relatively low, with IC_50_s reported in the micromolar range. This might complicate clinical development of MPS1-IN-3, as micromolar concentrations might be difficult to achieve in the brains of glioblastoma patients. The brain is well-protected by drug efflux transporters that are present at the blood-brain barrier (BBB) and restrict brain entry of exogenous compounds [11], even when BBB integrity is compromised [12, 13]. In fact, vincristine is one of many drugs that are efficiently transported by efflux transporters [14, 15], and multiple clinical trials investigating vincristine against glioblastoma have failed [16, 17]. In contrast, temozolomide is only a relatively weak substrate and is able to penetrate the brain at sufficient concentrations to achieve antitumor efficacy[18], and active against GBM in the clinic [1]. Thus, in order to show the potential of MPS1 inhibitors for treatment of GBM, preclinical studies should focus on highly potent compounds that have good BBB penetration, used either as monotherapy or in combination with the standard-of-care temozolomide chemo-radiotherapy.

A small-molecule MPS1 inhibitor (NTRC 0066-0) was developed that possesses superior potency compared to 11 other MPS1 inhibitors that were evaluated in parallel, including BAY 1217389, reversine, MPS1-IN-3 and NMS-P715 [19]. NTRC 0066-0 demonstrated markedly increased target residence time and the lowest IC_50_s of all MPS1 inhibitors against a panel of 66 cell lines. This makes NTRC 0066-0 a clear frontrunner among MPS1 inhibitors and a promising candidate to be tested in preclinical models of GBM. We here report that NTRC 0066-0 exhibits low nanomolar potency against a panel of GBM cell lines *in vitro*. NTRC 0066-0 does not synergize with microtubule poisons or DNA damaging chemotherapeutics. Although we found that NTRC 0066-0 is a substrate of the efflux transporter P-glycoprotein (P-gp; ABCB1), it has a good brain penetration and good oral bioavailability in mice. We investigated the *in vivo* antitumor efficacy of NTRC0066-0 using several dosing schedules and combination therapy settings, namely: dosing NRTC0066-0 at the maximum tolerated dose (MTD) every other day and twice-daily; as monotherapy in treatment-naïve tumors and at recurrence following temozolomide chemo-radiotherapy; and in a neo-adjuvant setting combined with temozolomide and radiotherapy. Although signs of tumor growth delay were observed in individual mice receiving treatment in a recurrent GBM setting, NTRC 0066-0 could not improve survival of mice carrying orthotopic GBM tumors in any of the settings tested. These findings indicate that developing MPS1 inhibitors for treatment of GBM will be challenging and future studies should aim to identify a population of GBM patients that might benefit from MPS1 inhibitor therapy in the recurrent setting.

## 2. METHODS

### 2.1 Drugs

NTRC 0066-0 was synthesized at the Netherlands Translational Research Center B.V. (Oss, The Netherlands) according to the procedure described in de Man *et al*. [20]. Temozolomide was purchased from Sigma-Aldrich (St. Louis, MO) for *in vitro* assays and TEVA Pharma (Haarlem, The Netherlands) for *in vivo* experiments. Vincristine was obtained from Pfizer (New York, NY), docetaxel from Hospira (Lake Forest, IL) and hydroxyurea from Sigma. Elacridar was provided by GlaxoSmithKline (Research Triangle Park, NC) and zosuquidar by Eli Lilly (Indianapolis, IN).

### 2.2 Cell culture

The serum-cultured GBM cell lines U87 (RRID:CVCL_0022) and HS683 (RRID:CVCL_0844) were purchased from the ATCC (Manassas, VA). T98G (RRID:CVCL_0556) and LN-428 (RRID:CVCL_3959) cells were kindly provided by Dr. Conchita Vens (Netherlands Cancer Institute, Amsterdam, The Netherlands), E98 cells were obtained from Prof. Dr. William Leenders (Radboud University Medical Center, Nijmegen, The Netherlands), and LN-751 (RRID:CVCL_3964) were generously made available by Prof. Dr. Monika Hegi (Centre Hospitalier Universitaire Vaudois, Lausanne, Switzerland). GSC lines were isolated from murine tumors that were generated by stereotactic injection of Lenti–Cre vector in the striatum of *Egfr^vIII^;p15^Ink4b^/p16^Ink4aF/F^* (GSC750), *Egfr^vIII^;p15^Ink4b^/p16^Ink4aF/F^;Pten^F/F^* (GSC578) or *Egfr^vIII^;p15^Ink4b^/p16^Ink4aF/F^;Pten^F/F^;Tp53^F/F^* (GSC556) mice, as previously described [21].

All non-GSC lines were cultured in MEM supplemented with 10% FBS, 1% L-glutamine, 1% sodium pyruvate, 1% MEM vitamins, 1% non-essential amino acids and 1% penicillin/streptomycin (all from Life Technologies, Carlsbad, CA) under 37 °C and 5% CO_2_ conditions. GSC lines were cultured as neurospheres on ultra-low attachment plates (Corning Inc.; Corning, NY) in 50% Neurobasal medium and 50% DMEM/F12 + GlutaMAX supplemented with 1% penicillin/streptomycin, 2% B-27 minus vitamin A (all Life Technologies), 10 ng/ml EGF and 10 ng/ml bFGF (both PeproTech, London, UK).

### 2.3 Cell survival assays

Cells were plated in 24-wells plates at low cell densities (2,000–10,000 cells/well) and exposed to different drugs 24 hours later. Wells were pre-coated with poly-ornithine (15 µg/ml for 3 hours at room temperature) and subsequently with laminin (7.5 µg/ml for 1 hours at 37°C) prior to seeding GSCs to make them adherent without inducing differentiation. NTRC0066-0 was dissolved in DMSO (Sigma-Aldrich) for *in vitro* experiments and final DMSO concentrations never exceeded 0.1%. When untreated wells were confluent (approximately 6-10 days after seeding, dependent on the cell line), cells were fixed and stained using a solution containing 0.5% crystal violet (w/v; Sigma-Aldrich) and 6% glutaraldehyde (v/v; Honeywell, Morris Plains, NJ). Plates were imaged using a Chemi-Doc XRS+ (Bio-rad, Hercules, CA) and analyzed using the Colony Area plugin of ImageJ [22]. IC_50_ curves were fitted and plotted using the ‘log(inhibitor) vs. normalized response – Variable slope’ fitting procedure of GraphPad Prism v7 (GraphPad Software, La Jolla, CA). Combination indices (CI) were calculated according to the method described by Chou and Talalay, in which CI < synergism, CI = 1 additivity and CI > antagonism[23].

### 2.4 Animals

Mice were housed at 20.9 ᵒC on a 12 hour light/dark cycle with food and water *ad libitum*. All animal housing and studies were approved by the Animal Experiments Committee of the Netherlands Cancer Institute and conducted according to national law and institutional guidelines.

### 2.5 Pharmacokinetic studies

The pharmacokinetics of NTRC 0066-0 were analyzed in wild-type (WT), *Abcb1a/b^-/-^*, *Abcg2^-/-^*, *Abcb1a/b;Abcg2^-/-^* and *Abcb1a/b;Abcg2;Abcc4^-/-^* FVB mice. All knockout mice strains have been developed at the Netherlands Cancer Institute [24–27]. For i.v. administration, NTRC0066-0 was formulated in DMSO:Cremophor EL:saline (1:1:8 v/v) and injected at a dose of 5 mg/kg. For i.p and oral administration, NTRC0066-0 was administered at 5, 10 or 20 mg/kg as indicated. Serial blood sampling was done by tail vein bleeding at specific time points. At terminal time points, blood was drawn by cardiac puncture followed by tissue collection. Plasma was obtained from whole blood by centrifugation (5 min, 5,000 rpm, 4°C) and tissues were weighed and homogenized using a FastPrep®-24 (MP-Biomedicals, NY) in 1% (w/v) bovine serum albumin (Sigma-Aldrich) in water.

### 2.6 LC-MS/MS analysis

NTRC 0066-0 was extracted from *in vitro* and *in vivo* samples using liquid–liquid extraction with diethyl ether. Another MPS-1 inhibitor, Compound 5 (Cpd-5) [9], was added to the samples as an internal standard. The dried organic phase was reconstituted in 30% methanol in water (v/v) [28]. Samples were measured using an LC-MS/MS system comprising an UltiMate 3000 LC Systems (Dionex, Sunnyvale, CA) and an API 4000 mass spectrometer (Sciex, Framingham, MA). Samples were injected on to a Securityguard C18 pre-column (Phenomenex, Utrecht, The Netherlands) connected to a ZORBAX Extend-C18 column (Agilent, Santa Clara, CA). Elution was done at a flow rate of 0.2 mL/min using a 5 minute gradient from 20% to 95% B (mobile phase A was 0.1% HCOOH in water (v/v) and mobile phase B was methanol). 95% B was maintained for 3 min followed by re-equilibration at 20% B. Multiple reaction monitoring parameters were 566.2/391.4 (NTRC0066-0) and 581.6/397.0 (internal standard). Analyst^®^ 1.6.2 software (Sciex) was used for system control and data analysis.

### 2.7 Orthotopic xenograft studies

Orthotopic GBM tumors were induced in *Abcb1a/b;Abcg2^-/-^* mice by stereotactic injection of E98 cells expressing firefly luciferase (200,000 cells in 2 µL HBSS (Life Technologies) at an injection speed of 1 µL/min) at 2 mm lateral, 1 mm anterior and 3 mm ventral to the bregma. Tumor growth was monitored using bioluminescence imaging by injecting D-luciferin (150 mg/kg i.p.; Promega, Madison, WI) and subsequently imaging mice on an IVIS Spectrum Imaging System (PerkinElmer, Waltham, MA). Survival was monitored daily and the humane endpoint was defined as body weight loss exceeding 20%. Time-to-recurrence was defined as the number of days elapsed until the tumor regained its pre-treatment size. Tumors were treated with CT-guided radiotherapy using an X-RAD 225Cx system (Precision X-Ray, North Branford, CT), temozolomide (10 mg/kg p.o. freshly prepared from powder in an aqueous solution containing 10% v/v ethanol), NTRC0066-0 (20, 10 or 5 mg/kg p.o. in an aqueous solution containing 10% v/v DMSO and 10% v/v Cremophor EL (Life Technologies)) or different combinations thereof, as described per experiment in the Figure legends.

### 2.8 Quantification of mitotic errors

Tumor-bearing mouse brains were formalin-fixed, paraffin-embedded and coronal sections (10 µm) were sliced using an RM2255 microtome (Leica, Wetzlar, Germany). Sections were subsequently deparaffinized using xylene, incubated in citrate buffer for antigen retrieval, blocked with 4% (w/v) BSA in TBS-T, incubated with rabbit-α-pH3^Ser10^ (1:250 for 3 hours at room temperature; Merck Millipore, Burlington, MA), incubated with goat-α-rabbit-AF568 (1:250 for 45 minutes at room temperature; Thermo Fisher Scientific, Waltham, MA) and mounted with Vectashield hardset antifade mounting medium with DAPI (Vector Laboratories, Burlingame, CA). Mitotic errors were imaged and scored using a Deltavision ultra high resolution microscope (Applied Precision Inc., Issaquah, WA).

### 2.9 Magnetic resonance imaging

Magnetic resonance imaging was done using a sequence consisting of T2-weighted, T1-weighted pre-contrast and T1-weighted post-gadoterate meglumine (Dotarem^®^; Guerbet; Villepinte, France) contrast imaging on a BioSpec 70/20 USR (Bruker, Billerica, MA) system, as described previously [18].

### 2.10 Histology and Immunohistochemistry

Mouse heads were fixed in 4% (v/v) formaldehyde and 5% (v/v) glacial acetic acid overnight, and subsequently decalcified in 6.5 % (v/v) formic acid at 37 °C for 4 days. Decalcified tissues were paraffin embedded and cut into 4 µm coronal sections that were stained with hematoxylin and eosin (H&E), and for human Vimentin (1:4,000; M0725; DakoCytomation; Glostrup, Denmark), P-gp (1:200; 13978; Cell Signaling Technology; Danvers, MA) and BCRP (1:400; ab24115; Abcam; Cambridge, UK). Sections were imaged and processed using an Aperio AT2 system and ImageScope software (v12; both Leica).

### 2.11 Pharmacokinetic and statistical analysis

PK solver was used to determine pharmacokinetic parameters [29]. The standard error of the oral bioavailability was calculated using the formula below:

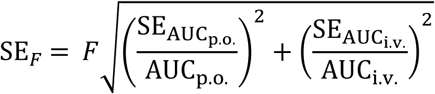

All comparisons involving more than two experimental groups were done using one-way analysis of variance followed by *post hoc* Bonferroni tests. Statistical differences between PK parameters were calculated as Bonferroni corrected p-values using multiple Student’s t-tests. Kaplan-Meier survival curves were analyzed using the log-rank test. Differences were considered statistically significant when *p* < 0.05.

## 3. RESULTS

### 3.1 NTRC0066-0 is cytotoxic against GBM cell lines with low nanomolar potency but does not synergize with chemotherapeutics

We first established the cytotoxic potency of NTRC 0066-0 against a panel of GBM cell lines *in vitro* using continuous drug exposure. The IC_50_ of NTRC 0066-0 against six serum-cultured cell lines was very uniform and in the low nanomolar range, varying from approximately 20 nM to 40 nM (Figure 1A). Moreover, the IC50 curves were very steep, with cell survival declining from 100% to 0% within one decade of concentration. These results were validated using three independent *Egfr^vIII^*-driven murine glioma stem cell (GSC) lines, as GSCs more accurately resemble human GBM than classical serum-cultured lines [30]. We observed similarly low nanomolar IC_50_s against *Egfr^vIII^;p15^Ink4b^/p16^Ink4a^*^-/-^ (GSC750), *Egfr^vIII^*;*p15^Ink4b^/p16^Ink4a^*^-/-^*;Pten^-/-^* (GSC578) and *Egfr^vIII^*;*p15^Ink4b^/p16^Ink4a^*^-/-^*;Pten^-/-^;Tp53^-/-^* (GSC556) GSCs, confirming the potent efficacy of NTRC0066-0 against GBM cells *in vitro* (Figure 1B).

**Figure 1.**
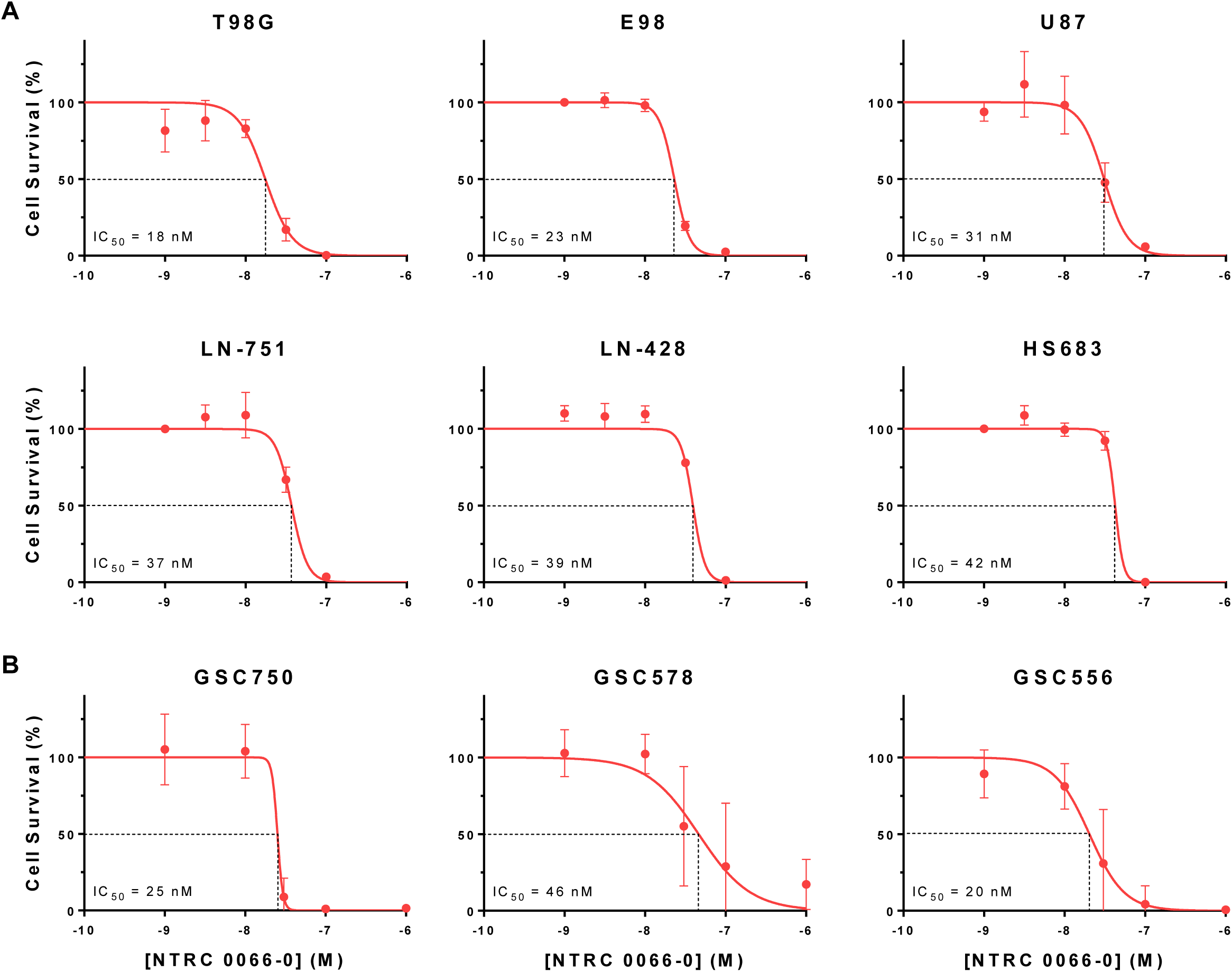
*In vitro* cytotoxicity of NTRC 0066-0 against a panel of GBM cell lines. (**A**) IC_50_ curves of NTRC0066-0 against six different serum-cultured GBM cell lines. Data are means ± SD; n ≥ 3. (**B**) IC_50_ curves of NTRC 0066-0 against *Egfr^vIII^;p15^Ink4b^/p16^Ink4a^*^-/-^ (GSC750), *Egfr^vIII^;p15^Ink4b^/p16^Ink4a^*^-/-^*;Pten^-/-^* (GSC578) and *Egfr^vIII^;p15^Ink4b^/p16^Ink4a^*^-/-^*;Pten^-/-^;Tp53^-/-^* (GSC556) glioma stem cell (GSC) lines. Data are means ± SD; n ≥ 8.

We next investigated whether NTRC 0066-0 can synergize with other chemotherapeutics to induce cytotoxicity in GBM cell lines. NTRC 0066-0 was combined with the microtubule poison vincristine (Figure 2A), the microtubule stabilizing agent docetaxel (Figure 2B), the alkylating agent temozolomide that is part of the standard-of-care treatment for GBM (Figure 2C), and the ribonucleotide reductase inhibitor hydroxyurea that induces replication stress (Figure 2D). NTRC0066-0 did not synergize with any of the tested chemotherapeutics and even displayed moderate to, in some instances, strong antagonism, as indicated by Combination Indices (CIs) over 1.0 up to 2.9 (Figure 2F). Although NTRC 0066-0 was continuously present in the previous experiments, exposure to 100 nM for about 8 h was already sufficient to reduce cell survival to about 25%. The addition of vincristine or docetaxel did not change the required exposure time (Figure 2E).

**Figure 2.**
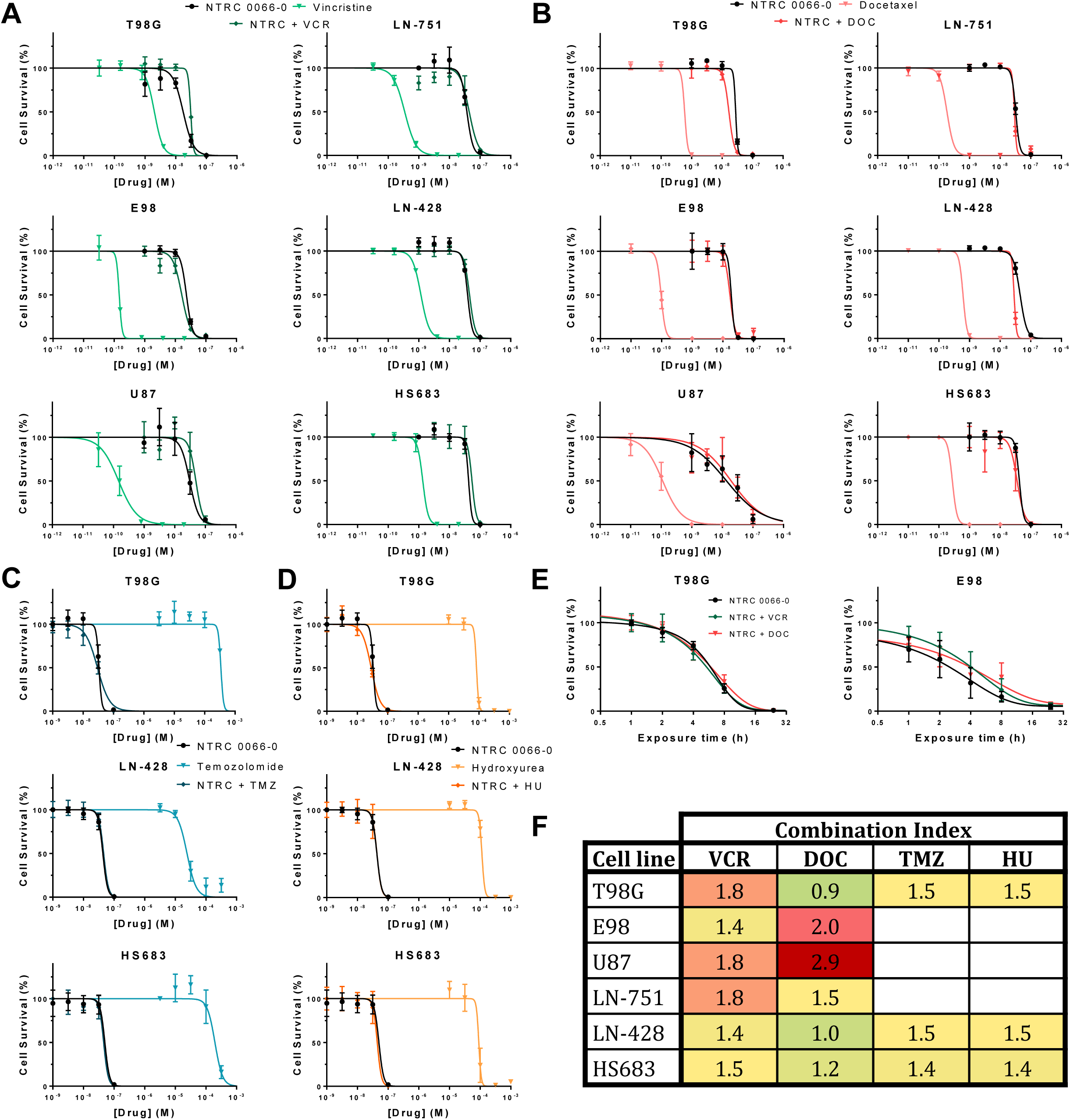
NTRC 0066-0 does not synergize with classic chemotherapeutics *in vitro*. IC_50_ curves of NTRC 0066-0 in combination with (**A**) vincristine (VCR; 0.2 nM), (**B**) docetaxel (DOC; 0.2 nM for LN-428 and T98G, 0.1 nM for HS683 and LN-751, and 0.05 nM for E98 and U87), (**C**) temozolomide (TMZ; 200 µM for T98G, 100 µM for HS683 and 10 µM for LN-428) and (**D**) hydroxyurea (HU; 50 µM) against different serum-cultured GBM cell lines. (**E**) Exposure time– response curves of NTRC 0066-0 (100 nM) alone or in combination with VCR or DOC against T98 (0.2 nM VCR or DOC) and E98 (0.2 nM VCR or 0.05 nM DOC) GBM cells. (**F**) Combination indices (CI) of drug combinations depicted in (A-D), as described by Chou and Talalay: CI < 1 depicts synergism, CI = 1 additivity and CI > antagonism [23]. Data are means ± SD; n = 4.

### 3.2 NTRC0066-0 achieves good brain penetration, despite being transported by P-gp in vitro and in vivo

ATP-binding cassette (ABC) efflux transporters are expressed at the BBB and limit the intracranial efficacy of anticancer agents by restricting their brain entry [11]. The most dominant ABC transporters at the BBB are P-gp (also known as MDR1 or ABCB1) and breast cancer resistance protein (BCRP; or ABCG2). Previously, it was shown that NTRC 0066-0 does not inhibit the activity of P-gp in a calcein-AM efflux assay [31]. Here, we determined whether NTRC 0066-0 is a substrate of ABC transporters at the blood-brain barrier. To study the impact of ABC transporters on the pharmacokinetics and brain penetration of NTRC 0066-0, we used mouse strains genetically lacking one or multiple transporters. We first assessed the impact of P-gp, BCRP and multi-drug resistance protein 4 (MRP4; ABCC4) on the brain penetration of NTRC 0066-0 at 1 h after i.v. administration of 5 mg/kg. Compared to FVB WT mice, *Abcb1a/b^-/-^* mice had markedly increased NTRC 0066-0 brain concentrations and brain-plasma ratios, while the plasma concentration was similar (Figure 3A). In contrast, no differences in plasma and brain concentrations were observed in *Abcg2^-/-^* mice. Furthermore, *Abcb1a/b;Abcg2^-/-^* and *Abcb1a/b;Abcg2;Abcc4^-/-^* mice did not display further increased NTRC 0066-0 brain concentration compared to *Abcb1a/b^-/-^* mice. Together, these data indicate that P-gp, but not BCRP and MRP4, limit the brain penetration of NTRC 0066-0. Importantly, NTRC 0066-0 displayed a high brain-plasma ratio (approximately 4) in WT mice despite being transported by P-gp, suggesting that it might be a useful candidate MPS1 inhibitor to test for treatment of GBM.

**Figure 3.**
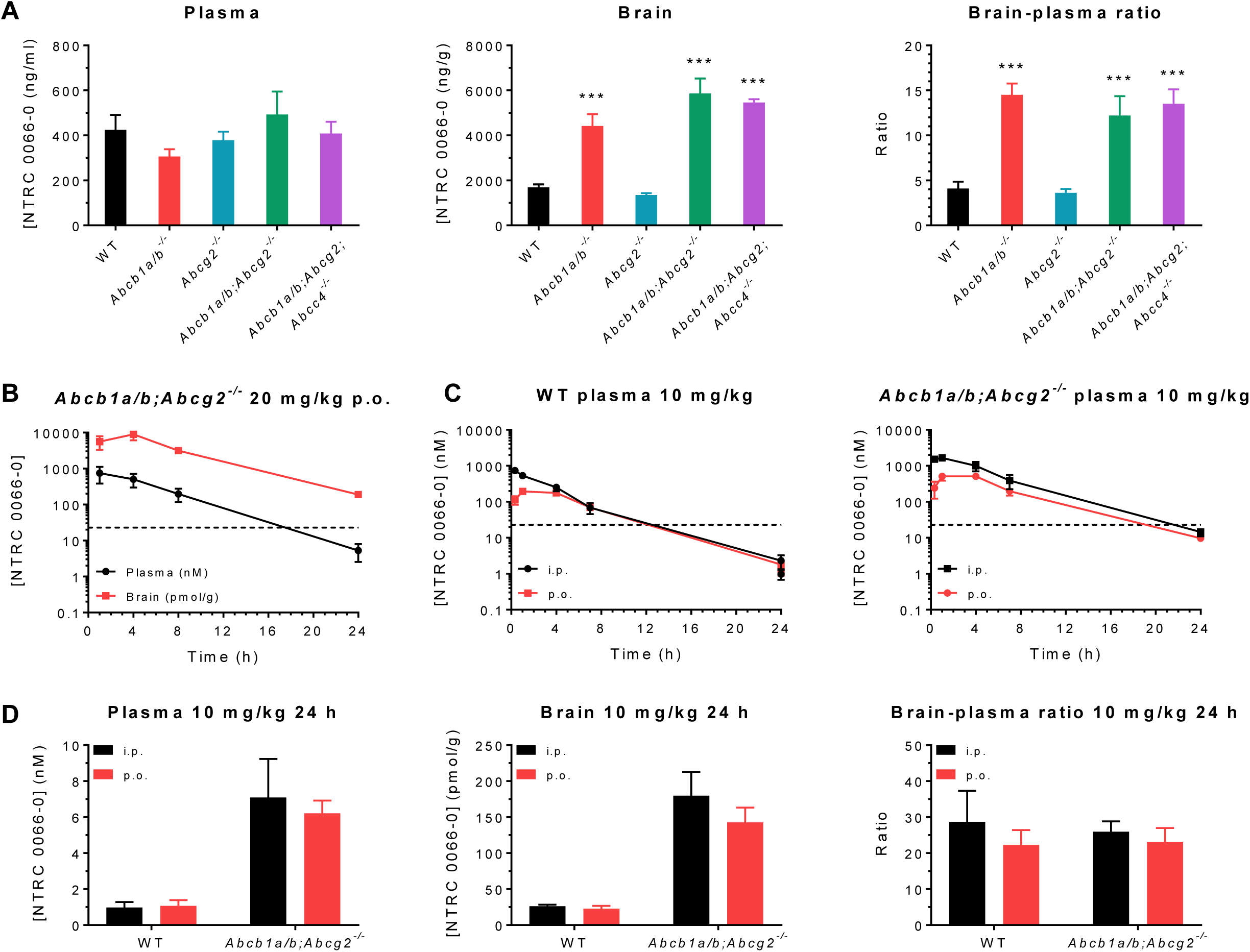
Impact of P-gp, BCRP and MRP4 on the pharmacokinetics of NTRC 0066-0. (**A**) Plasma concentration, brain concentration and brain–plasma ratio of NTRC 0066-0 following intravenous administration of 5 mg/kg to wild-type (WT), *Abcb1a/b^-/-^*, *Abcg2^-/-^*, *Abcb1a/b;Abcg2^-/-^* and *Abcb1a/b;Abcg2;Abcc4^-/-^* mice. (**B**) Plasma and brain concentration–time curves following administration of 20 mg/kg NTRC 0066-0 p.o. to WT mice. (**C**) Plasma concentration–time curves following administration of 10 mg/kg NTRC0066-0 p.o. and i.p. to WT and *Abcb1a/b;Abcg2^-/-^* mice. (**D**) Plasma concentration, brain concentration and brain–plasma ratio 24 hours after administration of 10 mg/kg NTRC 0066-0 p.o. and i.p. to WT and *Abcb1a/b;Abcg2^-/-^* mice. Data are mean ± SD; n ≥ 4; *** *p* < 0.001.

We next determined the oral bioavailability of NTRC 0066-0 in *Abcb1a/b;Abcg2^-/-^* and WT mice relative to i.p administered drug. We found that the plasma AUC of NTRC 0066-0 was about 2-fold higher both after i.p and oral dosing in *Abcb1a/b;Abcg2^-/-^* mice *vs.* WT mice, resulting in a similar oral bioavailability of about 50% in both strains (Figure 3C and Table 1). The terminal half-life of NTRC 0066-0 was significantly longer in *Abcb1a/b;Abcg2^-/-^* mice compared to WT mice (Figure 3C and Table 1). At 24 hours, we collected the brains of these mice. Elimination of NTRC 0066-0 from the brain was slower than from the systemic circulation, as the brain-to-plasma ratio was approximately 20 at 24 hours. Interestingly, the brain-to-plasma ratio was similar in both strains, although the concentration in *Abcb1a/b;Abcg2^-/-^* mice at 24 h was about 5-fold higher than in WT mice (Figure 3A, D).

**Table 1.** Pharmacokinetic parameters after oral and i.p. administration of NTRC 0066-0 to WT and *Abcb1a/b;Abcg2^-/-^* mice. AUC, area under the curve; *C*_max_, maximum concentration in plasma; *t*_max_, time to reach maximum plasma concentration; *t*_1/2_, plasma half-life; MRT, mean residence time; *F*, bioavailability. Data are represented as mean ± SD (n ≥ 4); ** *p* < 0.01, **** *p* < 0.0001.

|  | Parameter | Time (h) | Genotype |  |
| --- | --- | --- | --- | --- |
|  |  |  | WT | *Abcb1a/b;Abcg2*<sup>-/-</sup> |
| i.p.<br>10 mg/kg | Plasma AUC<sub>i.p.</sub> (nM·h) | 0-∞ | 2800 ± 450 | 11000 ± 2200*** |
|  | C<sub>max</sub> (nM) |  | 780 ± 130 | 1700 ± 160*** |
|  | t<sub>max</sub> (h) |  | 0.5 ± 0.3 | 0.8 ± 0.3 |
|  | t<sub>1/2</sub> (h) |  | 2.5 ± 0.2 | 3.4 ± 0.4* |
|  | MRT (h) |  | 3.2 ± 0.4 | 4.3 ± 0.4* |
| p.o.<br>10 mg/kg | Plasma AUC<sub>p.o.</sub> (nM·h) | 0-∞ | 1700 ± 150 | 4700 ± 710** |
|  | C<sub>max</sub> (nM) |  | 180 ± 6.7 | 550 ± 52*** |
|  | t<sub>max</sub> (h) |  | 1.8 ± 1.5 | 3.3 ± 1.5 |
|  | t<sub>1/2</sub> (h) |  | 2.8 ± 0.2 | 3.6 ± 0.2** |
|  | F (%) |  | 59 ± 11 | 43 ± 11 |
|  | MRT (h) |  | 4.7 ± 0.2 | 5.2 ± 0.5 |

In view of the higher NTRC 0066-0 levels in the absence of P-gp, we conducted the first proof-of-concept studies in *Abcb1a/b;Abcg2^-/-^* mice (see below). We first established that 20 mg/kg NTRC 0066-0 every other day for 11 days (*q.*2*d.*x11d) could safely be given orally to *Abcb1a/b;Abcg2^-/-^* mice. This schedule has been used previously in a breast cancer model in WT FVB mice [31]. Next, we determined the brain and plasma levels at 1, 4, 8 and 24 h after oral dosing of 20 mg/kg and found that the brain concentration remained well above the threshold level of *in vitro* efficacy (Figure 3B).

### 3.3 NTRC 0066-0 does not improve survival of mice carrying orthotopic GBM tumors, although signs of tumor growth delay are observed in a recurrent GBM model

To study the *in vivo* antitumor efficacy of NTRC 0066-0 in relevant GBM models, we orthotopically injected E98 cells, as this cell line was one of the most sensitive to NTRC 0066-0 *in vitro* (Figure 1A) and robustly engrafts in nude mice. Moreover, orthotopic E98 tumors are only slightly visible on gadolinium-enhanced magnetic resonance imaging (Figure 4A) and well-perfused by vessels expressing P-gp and BCRP (Figure 4B), indicating that the BBB in these tumors retains it functionality at least partly. We conducted all *in vivo* studies in *Abcb1a/b;Abcg2^-/-^* mice, as these have significantly higher NTRC 0066-0 exposure than WT mice (Figure 3) This allowed us to explore the concept of *in vivo* MPS1 inhibition therapy in the most optimal pharmacokinetic setting. Treatment was started when tumors were relatively small (bioluminescence signal of about 10^5^ photons per second; p/s) to allow a sufficient window for a 21 day treatment period. However, treatment of therapy-naïve E98 tumors with 20 mg/kg NTRC 0066-0 *q.*2*d.*x11d did not result in reduced tumor growth (Figure 5A) or improved survival (Figure 5B).

**Figure 4.**
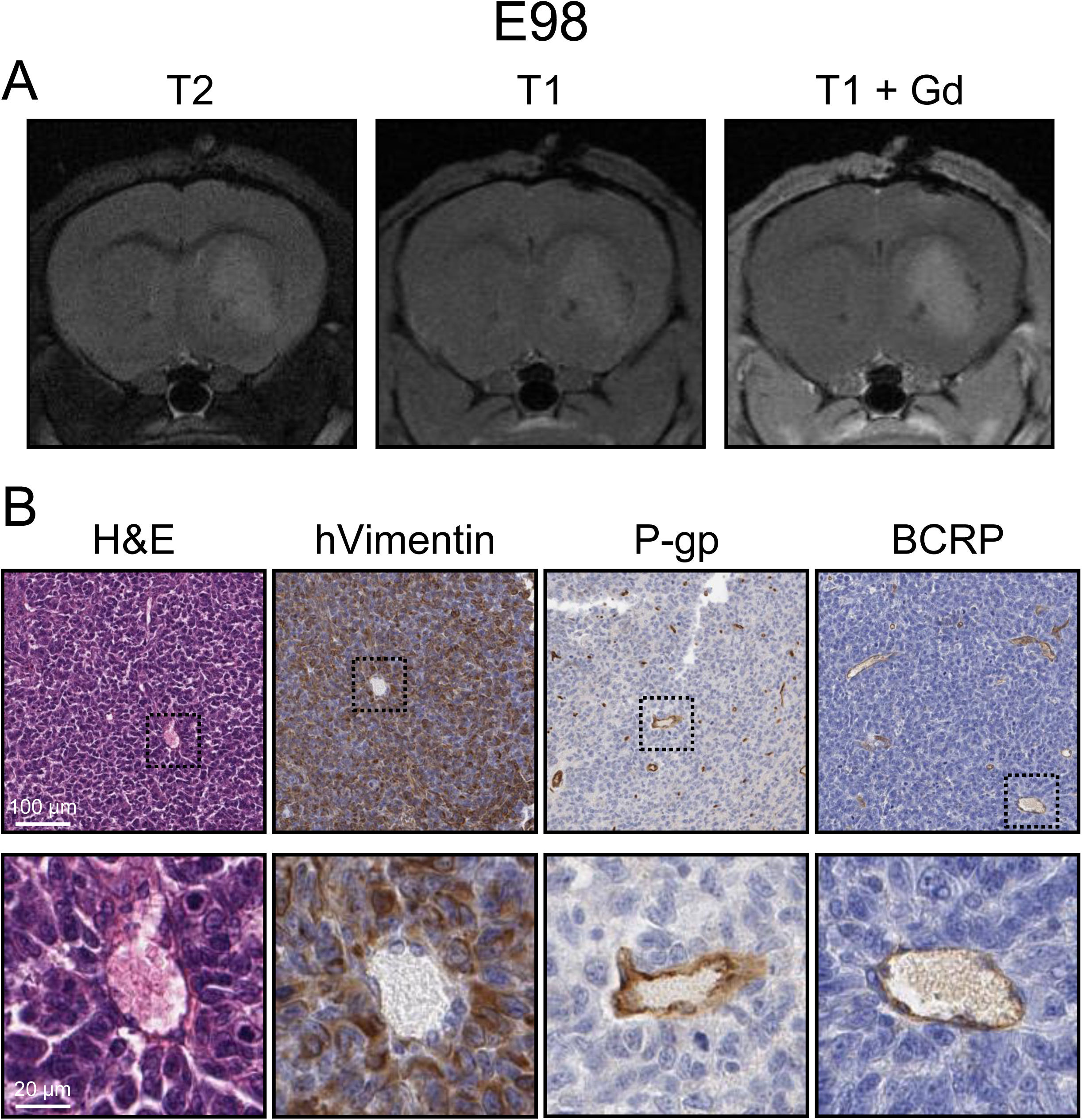
Characterization of the intracranial orthotopic E98 GBM model. (**A**) Magnetic resonance imaging of an orthotopic E98 tumor using a sequence consisting of T2-weighted, T1-weighted pre-contrast and T1-weight gadolinium contrast-enhanced (T1 + Gd) imaging. (**B**) Coronal E98 tumor sections stained for hematoxylin and eosin (H&E), human vimentin (hVimentin), P-glycoprotein (P-gp) and breast cancer resistance protein (BCRP).

**Figure 5.**
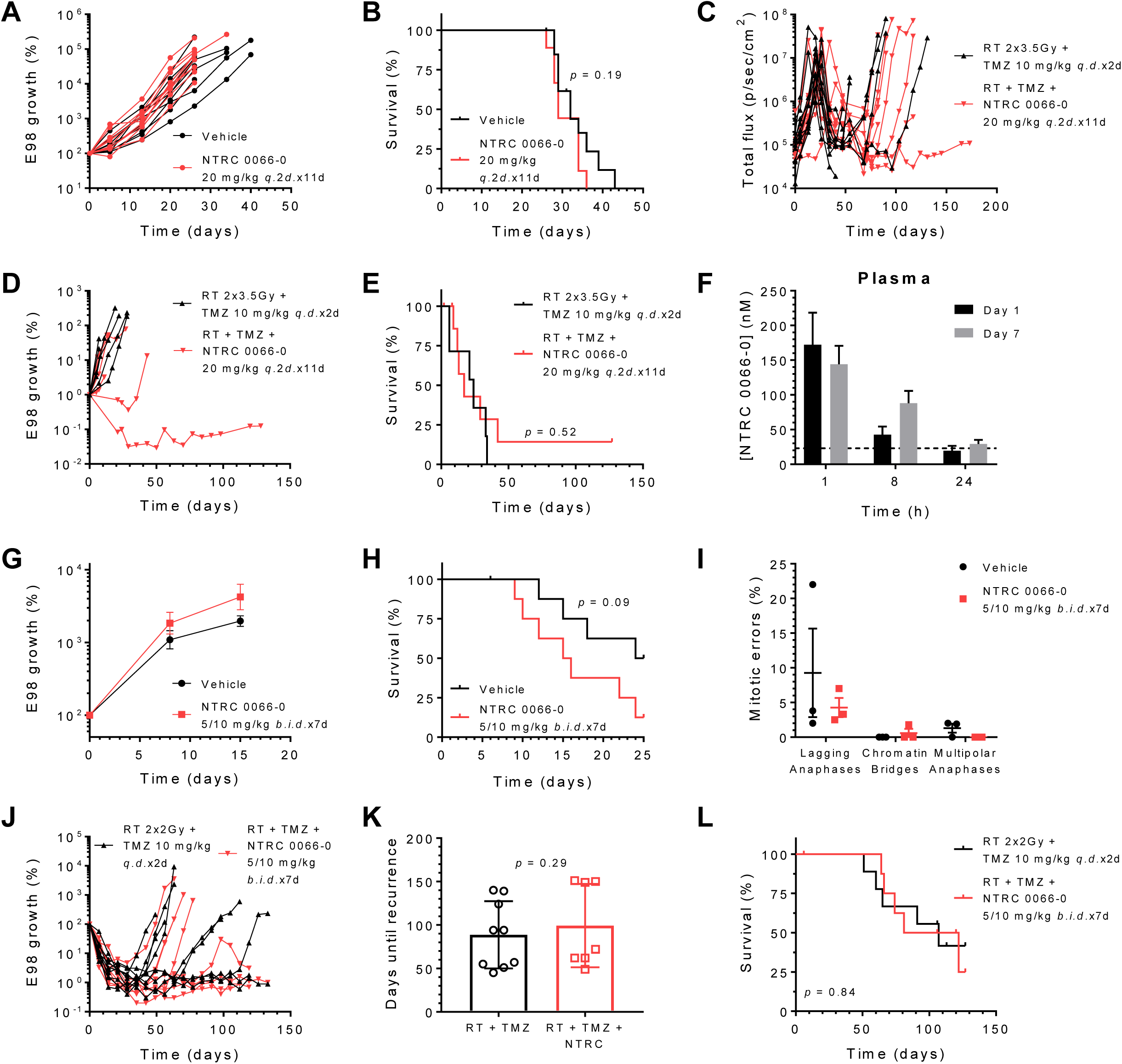
NTRC 0066-0 antitumor efficacy studies against orthotopic E98 GBMs in *Abcb1a/b;Abcg2^-/-^* mice. (**A**) Tumor growth and (**B**) survival of mice treated with NTRC 0066-0 administered at 20 mg/kg every other day for 11 days (*q.*2*d.*x11d). n = 10. (**C**) Tumor growth from tumor induction, (**D**) tumor growth stratified to time-of-recurrence and (**E**) survival from time-of-recurrence of mice treated with NTRC 0066-0 administered at 20 mg/kg *q.*2*d.*x11d when recurrence occurred after treatment with 2 consecutive days (*q.d.*x2d) of 3.5 Gy radiotherapy (RT) and 10 mg/kg temozolomide (TMZ) chemotherapy, or RT + TMZ alone. Treatment with RT + TMZ was started when tumor reached 10^7^ p/sec/cm^2^; treatment with NTRC 0066-0 was started when recurrence was detected. n = 7. (**F**) Plasma concentrations of NTRC 0066-0 in *Abcb1a/b;Abcg2^-/-^* mice at several time points of day 1 and day 7 of treatment with 5 mg/kg NTRC0066-0 in the morning and 10 mg/kg NTRC 0066-0 approximately 8 hours later for 7 consecutive days (5/10 mg/kg *b.i.d.*x7d). Data are mean ± SD; n = 5. (**G**) Tumor growth and (**H**) survival of mice treated with NTRC 0066-0 administered at 5/10 mg/kg *b.i.d.*x7d. Data are mean ± SE; n = 9. (**I**) Percentages of various classes of mitotic errors in end-stage tumors from (**G**). Data are mean ± SD; n ≥ 50 cells/tumor, n = 3 mice/group. (**J**) Tumor growth, (**K**) time-of-recurrence (**L**) survival of mice treated with 2 Gy RT and 10 mg/kg TMZ *q.d.*x2d followed by NTRC 0066-0 administered at 5/10 mg/kg *b.i.d.*x7d, or RT + TMZ alone. Data are mean ± SD; n = 9.

In the clinic, GBMs are treated with the DNA damaging therapies temozolomide (TMZ) and radiotherapy (RT), but uniformly recur. Since DNA damage is known to induce missegregations and aneuploidy [3], MPS1 is critical in protecting cells from mitotic cell death when encountering missegregations, and stable aneuploid cells are highly sensitive to MPS1 inhibition [33], we investigated whether NTRC 0066-0 could be effective against recurrent E98 tumors following TMZ + RT combination treatment (Figure 5C). TMZ + RT was given when tumors reached a size between 10^7^ and 10^8^ p/s and resulted in a substantial regression back to about 10^5^ p/s. Administration of 20 mg/kg NTRC 0066-0 (*q.*2*d.*x11d) was started at the time of recurrence and resulted in remarkable growth delay in 2 out of 7 animals (Figure 5D) with one long-term survivor (Figure 5E), whereas the other tumors progressed as rapidly as vehicle treated animals. The reason for this heterogeneity is unclear, but could be related to the type and extent of DNA damage that was induced by TMZ + RT treatment in individual mice. Furthermore, the lack of efficacy in 5 out of 7 mice might be attributed to insufficient brain exposure. The plasma concentration at 24 h after administration of 20 mg/kg was only 6 nM (Figure 3D) and although the brain penetration of NTRC0066-0 in *Abcb1a/b;Abcg2^-/-^* mice is 20-fold higher, the every other day (*q.*2*d.*) dosing schedule will result in very low levels in the period between 24 and 48 h, which may allow tumor tissue recovery in these drug-free periods.

In an effort to achieve more continuous exposure to NTRC 0066-0 in mice, we investigated a more dose-dense administration schedule. We found that we could safely administer 5 mg/kg p.o. in the morning, and 10 mg/kg p.o. approximately 8 hours later for 7 consecutive days. Based on our pharmacokinetic studies, the lowest plasma trough concentration of NTRC 0066-0, which is reached each morning just prior to the next dose, remained above 20 nM (Figure 5F). This concentration was effective against E98 cells *in vitro* and already higher than the plasma concentration observed 24 hours after administration of 20 mg/kg p.o. to *Abcb1a/b;Abcg2^-/-^* mice (Figure 1A and Figure 3B). Notably, the brain–plasma ratio is more than 20 in *Abcb1a/b;Abcg2^-/-^* mice. Treatment with this dose-dense schedule was started when the bioluminescence signal of the tumors was between 5–10. 10^6^ p/s. Unfortunately, however, even under these most optimal conditions, we again did not observe any delay in tumor growth (Figure 5G) or survival (Figure 5H) when E98 tumors were treated with NTRC 0066-0 monotherapy. In fact, treatment with NTRC 0066-0 appeared to accelerate tumor growth and shorten survival. Although these effects were not statistically significant, this trend was seen in two independent experiments (Figure 5A-B; *p* = 0.19) and (Figure 5G-H; *p* = 0.09) In line with this lack of antitumor efficacy, NTRC0066-0 did not increase the number of mitotic errors in these tumors (Figure 5I). Notably, we only observed some cases of lagging anaphases, but very few chromatin bridges and multipolar anaphases.

In the same series, we also studied the dense NTRC 0066-0 dosing schedule (5/10 mg/kg *b.i.d.*x7d) in combination with chemo-radiotherapy. The recurrent tumor setting as performed above was not a feasible option, because a sufficiently powered study may require up to 40 animals in the treatment cohort. We therefore tested it in an adjuvant setting where we started NTRC 0066-0 on day 3 following two days of RT + TMZ (2 Gy + 10 mg/kg *q.d.*x2d). In this setting, we again could not observe any effects on tumor growth (Figure 5J), as the time-to-recurrence was not different between treatment groups (Figure 5K). Concordantly, survival was also not affected (Figure 5L). In summary, NTRC 0066-0 did not improve survival of mice carrying orthotopic GBM tumors in any of the settings tested., although signs of tumor growth delay were observed in 2 out of 7 (30%) mice treated in the recurrent GBM setting (Figure 5D-E).

## 4. DISCUSSION

This study explored the potential of MPS1 inhibition as a therapeutic strategy for treatment of GBM, a devastating primary brain tumor. The MPS1 inhibitor NTRC 0066-0 efficiently induced cytotoxicity in multiple GBM and GSC cell lines with low nanomolar potency, obviously demonstrating the intrinsic potential of MPS1 inhibition to induce cytotoxicity in GBM cells. Moreover, the BBB penetration of NTRC 0066-0 is high, allowing good distribution throughout the brain. However, there was no good translation to *in vivo* antitumor efficacy, as we did not observe any robust tumor growth delay or improved survival, even in the most optimized pharmacokinetic settings using *Abcb1a/b;Abcg2^-/-^* mice and the highest tolerable dose-dense oral administration schedule. In contrast, other studies have reported promising efficacy of MPS1 inhibition in preclinical tumor models [5, 8, 9, 31, 34–37]. Single compound efficacy of NTRC 0066-0 was shown in orthotopic models of the triple-negative breast cancer cell line MDA-MB-231 [31, 38] and a subcutaneous model of the lung cancer cell line A427 [38]. Several observations may help to explain the lack of efficacy of NTRC 0066-0 in GBM models, and understanding these may enable translation of the *in vitro* potential of MPS1 inhibition to *in vivo* treatment of GBM.

First, *in vivo* antitumor efficacy of MPS1 inhibition as monotherapy is rarely reported. In most studies, efficacy is only observed when MPS1 inhibitors are combined with classic chemotherapeutics that target microtubules such as vincristine [5], docetaxel [9, 31] or paclitaxel [8, 37]. Combination therapy with taxane drugs has only been studied in extracranial tumor models. However, such combinations are not attractive for treatment of GBM, as taxanes have a poor brain penetration as a result of efficient efflux by P-gp at the BBB [39, 40]. Although vincristine combination therapy has shown some modest efficacy in preclinical GBM models, vincristine is also a P-gp substrate [15, 41] and is clinically ineffective against GBM [16, 17]. Combining MPS1 inhibitors with newer microtubule stabilizers that have improved brain penetration might offer a more promising perspective [42], albeit one that requires further investigation.

Second, most studies that report monotherapy efficacy of MPS1 inhibition used subcutaneous models of extracranially occurring cancers such as colorectal cancer, cervical cancer and triple-negative breast cancer [8, 31, 35–38]. There are several potentially important differences between subcutaneous tumor models and the brain tumor model that we used. Within the brain microenvironment, E98 cells need about 7-10 weeks to develop to a size of about 30 to 50 cubic mm. Although this size is large enough to kill the animal, it is considerably smaller than the 2000 cubic mm size that the subcutaneously growing tumors can reach in a much shorter period [31, 38]. The more rapid expansion of those tumors may cause a higher replication stress in tumor cells. Moreover, due to their rapid expansion with profound angiogenesis, subcutaneous cancer models are known to have a relatively aberrant vasculature [43]. This leaky vasculature facilitates intratumoral drug distribution and retention, leading to more continuous and relatively high local drug exposure. In fact, pharmacokinetic analysis of intratumoral drug concentrations show that MPS1 inhibitors can accumulate in subcutaneous tumors (unpublished data). The more long-term exposure could be one of the factors explaining the observed monotherapy efficacy in the subcutaneous setting. Such vascular artifacts of subcutaneous cancer models may not occur in orthotopic GBM models. Brain tumor models such as E98 grow relatively slow without evidence of necrosis as they are well perfused by vessels expressing BBB markers (Figure 4). Therefore, the drug concentration in the brain tumor likely follows the course of the drug level in normal brain. In the case of the intermittent (every other day) dose schedule this may result in considerable intervals of inadequate drug levels. Notably, however, when we changed to *b.i.d.* dosing, the efficacy did not improve. Instead, the trend towards a more rapid tumor progression that was already notable in the intermittent dose schedule (Figure 5A-B) became more pronounced with *b.i.d.* dosing (Figure 5G-H). This could imply that exposure to NTRC 0066-0 enables faster cycling of tumor cells by interference with checkpoint control, but without producing sufficient mitotic errors for causing cell death. In general, the level of mitotic errors that we observed in E98 tumors was low, as the majority of the tumors harbored less than 5% of mitotic errors. Since we analyzed only end-stage tumors, it could be that the absence of highly aberrant cells reflects the clearance from the general tumor population as a result of mitotic cell death. However, one would expect to observe a delayed tumor growth during NTRC 0066-0 treatment if the population encountering mitotic errors and subsequent cell death was substantial. Since we could not find such an effect, it is more likely that NTRC0066-0 5/10 mg/kg *b.i.d.*x7d did not induce mitotic errors in E98 cells in intracranial xenografts *in vivo*. If the induction of mitotic errors is more abundant *in vitro*, this might offer another explanation for the observed disconnect between the *in vitro* and *in vivo* efficacy of NTRC 0066-0 against these GBM cells. Although the intracranial E98 xenograft propagates steadily (Figure 5A, G), the doubling time of these tumors is longer than cells cultured *in vitro*. Culturing in the presence of ample amounts of nutrients, growth factors and high oxygen levels may result in more pronounced replication stress due to higher intrinsic DNA damage levels and more mitotic segregation errors than occurs in the *in vivo* context. Under such conditions, GBM cells may depend more on proper control of cell cycle checkpoints. If so, this might also explain why we found some indications of efficacy when NTRC 0066-0 was given when E98 tumors recurred after RT + TMZ treatment, as these modalities are DNA damaging agents that are known to induce mitotic errors. Signs of efficacy were observed, even when NTRC 0066-0 was administered using the intermittent dosing scheme of 20 mg/kg *q.*2*d.*x11d. Using this scheme, clear antitumor responses were observed in 2 out of 7 mice. The level of heterogeneity in therapy response is striking, and might be related to substantial differences in intrinsic levels of mitotic errors - especially lagging anaphases - between individual tumors that were observed even in untreated tumors (Figure 5I). On the one hand, understanding the requirement of this intrinsic heterogeneity might help to select a more optimal preclinical model to demonstrate the potential of MPS1 inhibition for treatment of GBM and could potentially contribute to identifying a patient group that is most likely to benefit from therapeutic strategies that involve MPS1 inhibition. On the other hand, the finding that MPS1 inhibition may cause an accelerated proliferation of a subpopulation of tumor cells is not an attractive prospect for further clinical development.

In summary, we observed profound cytotoxicity in GBM cells by the MPS1 inhibitor NTRC 0066-0 *in vitro*, but could not translate this finding to antitumor efficacy *in vivo*, despite a high brain penetration of NTRC 0066-0 and using the most dose-dense oral administration schedule that was tolerable. These data indicate that developing MPS1 inhibitors for treatment of GBM will be challenging.

